# A coarse-grained MD model for disorder-to-order transitions in polyQ aggregation

**DOI:** 10.1101/2025.03.06.641960

**Authors:** Maurice Dekker, Mark L. van der Klok, Erik Van der Giessen, Patrick R. Onck

**Affiliations:** Zernike Institute for Advanced Materials, University of Groningen, Groningen, The Netherlands

## Abstract

Polyglutamine (polyQ) aggregation plays a central role in several neurodegenerative diseases, including Huntington’s disease. To investigate the underlying mechanisms of polyQ aggregation, we developed a coarse-grained molecular dynamics model calibrated using atomistic simulations and experimental data. To assess the model’s predictive power beyond the calibrated parameter set, we systematically varied side chain hydrophobicity and hydrogen bonding strength to explore a broader range of aggregation pathways. These pathways ranged from nucleated growth to liquid-to-solid phase transitions. Through seeded aggregation simulations, we observed that amyloid growth occurs primarily in the β-sheet elongation direction, although growth through steric zippering was also observed. Longer polyQ sequences (Q48) exhibited significantly faster growth compared to shorter sequences (Q23), underscoring the role of chain length in aggregation kinetics. Our model provides a versatile framework for studying polyQ aggregation and offers a foundation for investigating broader aggregation mechanisms and sequence variations.

## Introduction

Polyglutamine (polyQ) aggregation is a key pathological characteristic of several neurodegenerative disorders, including Huntington’s disease, spinocerebellar ataxias, and other related conditions [1– 3]. These diseases are characterized by the expansion of trinucleotide repeats within specific genes, leading to proteins with abnormally long polyQ sequences [4, 5]. The length of these sequences is directly correlated with the onset and severity of the disease [6, 7], as longer polyQ tracts are more prone to aggregation into insoluble amyloid fibrils [8, 9]. These fibrils are believed to disrupt cellular functions and contribute to neuronal death [1, 10]. Despite the critical role of polyQ aggregation in neurodegeneration, the precise molecular mechanisms underlying polyQ aggregation remain largely elusive.

Two primary mechanisms have been proposed for the formation of insoluble amyloid fibrils from soluble monomers. The first is nucleated growth [9, 11], which posits that aggregation begins with a small, stable nucleus that serves as a template for the further addition of monomers, leading to fibril growth. The nucleation event is often considered the rate-limiting step in the aggregation process. The second mechanism involves a liquid-to-solid phase transition (LSPT) [12, 13], where polyQ monomers first undergo liquid-liquid phase separation, forming dense liquid-like droplets. Within these droplets, intrinsically disordered molecules gradually organize into a solid β-sheet-rich structure, ultimately resulting in mature amyloid fibrils. Both mechanisms underscore the complex interplay of protein concentration, molecular interactions, and environmental factors in driving polyQ aggregation.

Understanding the aggregation process of polyQ remains a significant challenge due to its inherent complexity and dynamic nature. PolyQ aggregation involves a series of poorly understood and transient intermediate states [14], including small oligomers and larger protofibrils, which are often short-lived and difficult to capture experimentally. Additionally, the aggregation pathway can vary significantly depending on factors such as concentration [11, 14, 15], temperature [16, 17] and the presence of molecular chaperones [18–20].

Molecular dynamics (MD) simulations have emerged as a powerful tool for exploring the behavior of proteins at the atomic level, offering insights into processes like protein folding and aggregation. However, the computational cost of all-atom simulations limits their use for studying large systems or long timescales [21], both of which are crucial to understanding the full scope of protein aggregation. To address this, coarse-grained (CG) models have been developed, simplifying the system by representing groups of atoms as single interaction sites. This significantly reduces computational demand, enabling simulations of larger systems on biologically relevant timescales while retaining essential aggregation features.

Current CGMD models [22–24], which commonly represent each amino acid as a single bead, have been very effective in studying phase separation [25–29]. Despite their success, these models have notable limitations when applied to the study of protein aggregation into ordered structures such as amyloid fibrils [30]. In particular, many CG models are unable to balance side chain interactions and backbone hydrogen bonding, which are crucial for fibril stability. These limitations hinder their ability to fully describe the transition from disordered aggregates to stable, fibrillar structures, highlighting the need for a model that explicitly accounts for these key interactions.

Polyglutamine presents an ideal candidate for the development of such a CG residue-scale model, as its sequence homogeneity simplifies parameterization—only glutamine interactions need to be optimized. Moreover, the well-documented relationship between polyQ length and aggregation propensity [6– 9] provides a clear benchmark for model validation. Several CG models have been developed to study polyQ aggregation, offering insights into early-stage aggregation and β-sheet formation [31–35]. However, these studies were generally unable to fully capture the transition from monomeric polyQ to mature amyloid fibrils. Many existing CG models incorporate both hydrogen bonding and side chain interactions but primarily promote β-sheet formation, failing to capture the full aggregation process into amyloid fibrils. In particular, they often neglect structural features essential for fibril stability, such as side chain zippering, which ensures the tightly packed, highly ordered architecture of mature amyloids [36]. As long as residue-scale models do not resolve how these interactions contribute to aggregation, they remain limited in their ability to reproduce experimentally observed stable aggregates.

In this article, we present a novel CGMD force field specifically designed to study polyQ aggregation.

This force field, referred to as the 2BPA-Q model, extends our previous CGMD model [22, 37] by incorporating specific features tailored to capture the characteristic behavior of polyQ sequences. We first describe the development of this force field, which involved several steps, including the calibration of hydrophobic interactions, hydrogen bonding, and side chain parameters based on experimental data and all-atom simulations. To systematically explore the aggregation pathways predicted by the model, we then construct a phase diagram by varying side chain hydrophobicity (λ_SC_) and hydrogen bonding strength (*ε*_HB_), allowing us to assess the model’s predictive power beyond the calibrated parameter set. Finally, we apply the fully parameterized model to seeded aggregation simulations, providing insight into the molecular mechanisms of polyQ amyloid growth. Through systematic optimization, we have created a model that accurately reproduces the known structural characteristics of both disordered polyQ monomers and structured amyloids, and provides valuable insights into the disorder-to-order transitions that drive polyQ aggregation. By bridging the gap between experimental observations and molecular-level understanding, the 2BPA-Q model opens the possibility to significantly advance our knowledge of protein aggregation and inform the development of new therapeutic strategies for polyQ-based neurodegenerative diseases.

## Methods

In this article, we present a two-bead-per-amino-acid (2BPA) model for polyQ aggregation simulations. This model is an extension of an earlier 1BPA (one-bead-per-amino-acid) model that represents each amino acid by a single coarse-grained bead positioned at the backbone alpha-carbon. The 1BPA model features backbone interactions for intrinsically disordered proteins (IDPs) extracted from Ramachandran data of coiled peptide fragments [37] and has been used to study transport through the nuclear pore complex [22, 38–40] and liquid-liquid phase separation [25, 28].

The 1BPA framework is used as a starting point for the CG model of polyQ presented in this work. The residue-specific bending and torsion interactions from the 1BPA force field [37] are adopted for the 2BPA backbone beads, and the potential form of the hydrophobic interaction is transferred to the 2BPA force field as well. A more detailed representation of the glutamine residue is obtained by introducing a second bead that represents the side chain and by implementing an interaction scheme to model the backbone hydrogen bonding interactions between glutamine residues (Fig. 1).

**Fig. 1.**
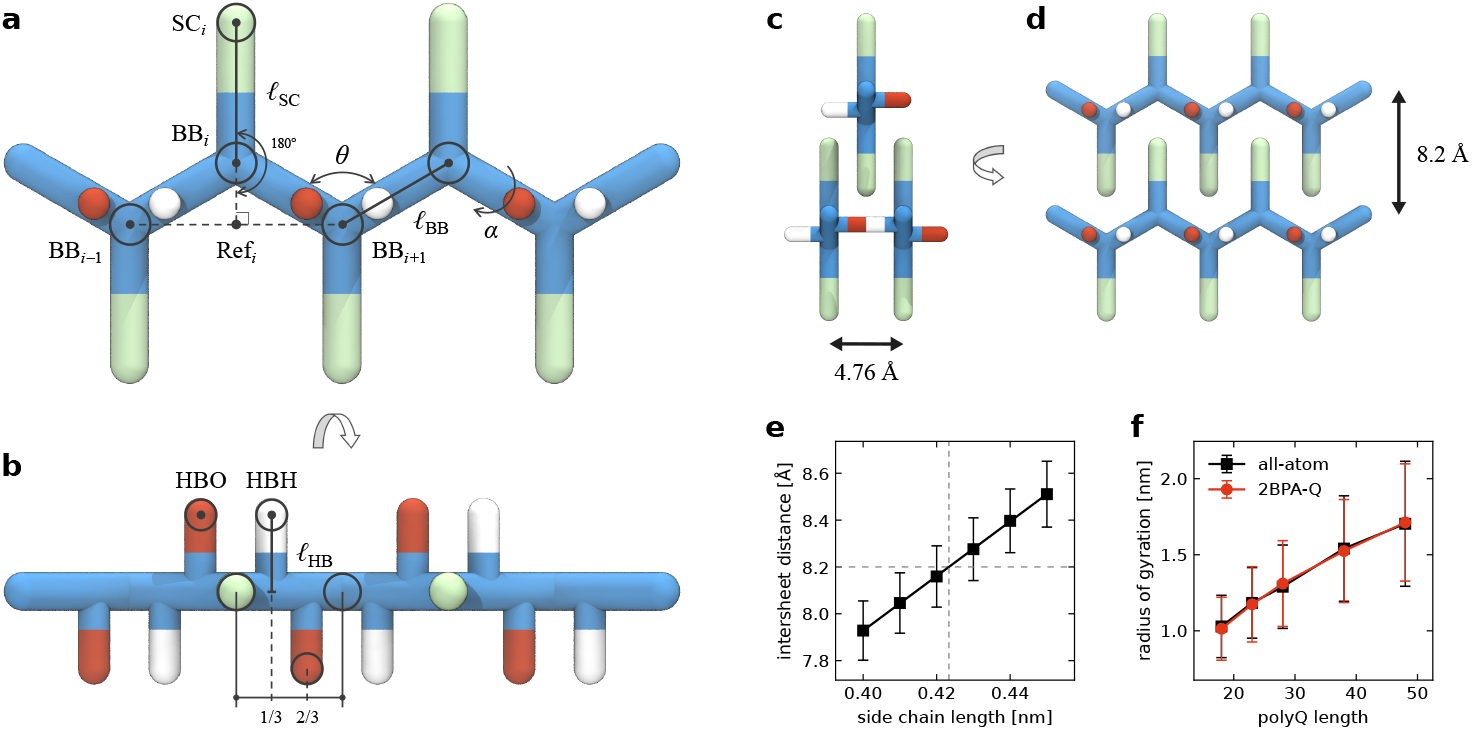
2BPA model for polyglutamine. (**a**) Coarse-grained representation of a polyQ5 chain. The stiffness of the protein backbone (BB, blue) is modeled accurately through pseudo-bending and torsion angles. Side chains (SC, green) are kept in place using a stiff 180^°^ bending potential to a virtual reference point (Ref) located in the middle of the previous and the next BB bead. An additional BB bead is added on each terminus to allow for the correct placement of the SC beads. (**b**) The hydrogen bonding beads (HBO, red; HBH, white) are placed perpendicular to the side chains at a distance *ℓ*_HB_ from the backbone bone bond. (**c**) The hydrogen bond length (*ℓ*_HB_ = 0.238 nm) is chosen such that the distance between two β-strands is 4.76 Å. (**d**) The distance between two β-sheets in a polyQ amyloid is 8.2 Å. (**e**) Intersheet distance in polyQ amyloids as a function of side chain length. The side chain length (*ℓ*_SC_ = 0.424 nm) is chosen such that the distance between β-sheets in polyQ amyloids matches the experimental intersheet distance of 8.2 Å [15, 41]. (**f**) Radius of gyration of single polyQ chains in all-atom simulations (black) and in the 2BPA-Q model (red). The optimal combination of backbone and side chain hydrophobicities (λ_BB_, λ_SC_) are determined such that the radius of gyration for various polyQ lengths matches with those obtained from all-atom simulations.

The bonded potential used for the backbone beads is taken directly from the 1BPA model [37], and is given by

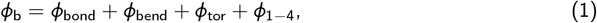

where the terms on the right-hand side are respectively the bond stretching, bending, torsion and 1–4 coupling potentials. The bond stretching between two covalently bonded backbone beads is governed by a stiff harmonic potential with an equilibrium distance *ℓ*_BB_ = 0.38 nm and force constant of 8038 kJ*=*nm^2^*=*mol. The bending and torsion potentials are obtained from Ramachandran data of coil regions of proteins, which were mapped to (*θ, α*)-space and converted into sequence-dependent potentials by Boltzmann inversion. The 1–4 coupling potential ensures proper sampling in (*θ, α*)-space because of the uncoupled backbone dihedrals. The reader is referred to Ghavami et al. [37] for a detailed description of the potentials in Eq. 1.

Hydrophobic interactions between backbone (BB) and side chain (SC) beads are described by a shifted 8–6 Lennard-Jones potential:

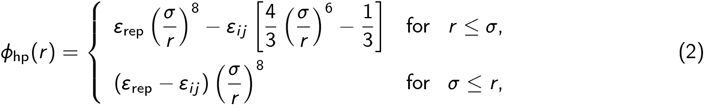

where the bead diameter is set to *σ* = 0.476 nm, ensuring that the combined volume of the BB and SC beads approximates the volume of a 1BPA bead [22, 28]. Furthermore, the diameter of the beads is the same as the strand separation in β-sheets [41, 42], allowing 2BPA peptides to form β-sheets without a steric clash. The interaction strength for each pair of beads (*i*, *j*) is given by the combination rule

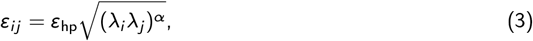

where λ_*i*_ is the relative hydrophobicity scale of bead *i* and the exponent *α* = 0.27 and the variables

*ε*_hp_ = 13 kJ*=*mol and *ε*_rep_ = 10 kJ*=*mol are taken from the 1BPA model [22].

### Side chain implementation

The implementation of side chain interactions can be done in multiple ways [43]. Here, we adopt a side chain construction inspired by the 3FD virtual site definition in GROMACS [44]. Early versions of our force field represented SC beads as virtual sites, meaning that their positions were determined relative to the BB beads rather than explicitly integrated. However, because these virtual sites were relatively distant from the backbone, the simulations became unstable once aggregation (i.e. phase separation or amyloid fibril formation) occurred. While reducing the integration time step to 0.01 ps could stabilize the system, we opted for a more robust alternative: treating side chain beads as massive particles with explicit integration.

To maintain the position of these massive side chain (SC) beads, we introduce a reference particle (Ref). This particle does not participate in nonbonded interactions; rather, it serves as a structural anchor to define the bonded interactions of the side chain. Specifically, for each SC bead, the reference point is positioned halfway along the line through BB beads *i* − 1 and *i* + 1 (see Fig. 1a):

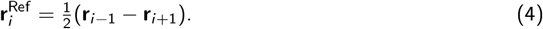

The position of the SC bead is restrained by a harmonic angle potential, with an equilibrium angle of 180^°^ and a force constant 1000 kJ*=*mol*=*rad^2^, between the SC, BB, and Ref beads. This approach ensures stability while preserving the intended structural constraints. Note that this construction method does not allow for the placement of side chains on the terminal residues. Therefore, an additional BB bead is placed on each terminus.

The glutamine side chain distance was calibrated based on experimental measurements of polyQ amyloid dimensions. In polyQ amyloids, the separation between two β-strands is 4.76 Å, while the distance between two β-sheets is 8.2 Å [15, 41], as illustrated in Fig. 1c,d. To determine the optimal side chain length, *ℓ*_SC_, we simulated 8×8 polyQ amyloid segments (i.e. 8 sheets of 8 strands) composed of polyQ32 strands with varying side chain lengths. These simulations revealed a linear relationship between intersheet distance and side chain length (Fig. 1e), leading us to identify the optimal value as *ℓ*_SC_ = 0.424 nm. Notably, these simulations were found to be insensitive to small variations in the hydrophobicity values of the BB and SC beads. Furthermore, we verified that, at the calibrated hydrophobicities, the optimal side chain length remains unchanged.

### Hydrogen bonding scheme

We adopt a simple but effective hydrogen bonding implementation similar to the CG model of Chen and Imamura [45, 46]. Backbone hydrogen bonds are modeled by explicitly including two hydrogen bonding (HB) beads for each residue, representing either a hydrogen (HBH; donor) or an oxygen (HBO; acceptor). These HB beads are virtual interaction centers, and their positions are calculated based on the positions of the BB beads, **r**_*i*_, using the following relations:

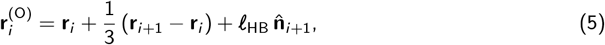

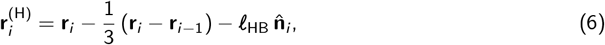

where

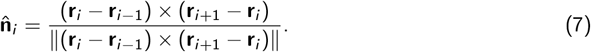

Here, the distance *ℓ*_HB_ is the perpendicular distance of the HB bead from the backbone bond (Fig. 1b), which is set to *ℓ*_HB_ = 0.238 nm in order to have the characteristic backbone-to-backbone distances of 4.76 Å between neighboring β-strands [41, 42, 47]. The HB beads are placed at 1/3 and 2/3 of the backbone bond, respectively, to make right-handed helices more stable with respect to left-handed helices [45] and results in a difference in stability between parallel and anti-parallel β-sheets. The construction of the HB interaction centers is realized in GROMACS [44] with a modified 3OUT virtual site definition (see Supplementary Methods for more details).

HBH beads do not interact with each other, nor do HBO beads. The interaction between HBO and HBH beads is described by a shifted Lennard-Jones potential:

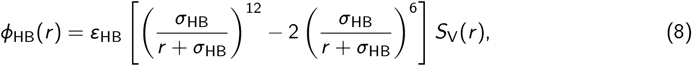

where *r* is the distance between HBO and HBH and *σ*_HB_ is a measure of the range of the interaction. An ideal hydrogen bond is formed when *r* = 0, resulting in a bond energy of *ε*_HB_. Based on values for β-sheets [48], we set the default hydrogen bonding strength to *ε*_HB_ = 6.6 kJ*=*mol. In accordance with the model of Chen and Imamura [45], the range of the hydrogen bonding interaction is set to *σ*_HB_ = 0.42 nm. To give the variable *σ*_HB_ a proper physical meaning, the hydrogen bonding potential is multiplied by the potential-switch function [49]:

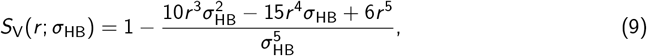

such that the potential is zero at *r* = *σ*_HB_. This hydrogen bonding potential is illustrated in Suppl. Fig. S1.

### Parametrization of the 2BPA-Q model

#### All-atom simulations

The hydrophobic interaction strengths of the BB and SC beads were calibrated using data from atomistic simulations of monomeric polyQ chains of various lengths. All-atom molecular dynamics simulations were performed using the GROMACS [44] software package (version 2021.5). Two distinct force fields were used to simulate the single-molecule systems: the amber99SB-disp (a99SB-disp) force field [50] with the modified TIP4P-D water model and the CHARMM36m (C36m) force field [51] with the TIP3P water model. Equations of motion were integrated with the leap-frog algorithm with a time step of 2 fs. For both force fields, the temperature was maintained at 300 K using the stochastic velocity rescaling (v-rescale) thermostat [52], with a coupling constant of 1 ps for a99SB-disp [50] and 0.1 ps for C36m [51]. The Parrinello–Rahman barostat [53] was utilized to maintain a pressure of 1 bar, with a coupling constant of 2.0 ps for a99SB-disp [50] and 1.0 ps for C36m [51], and an isothermal compressibility of 4.5 × 10^−5^ bar^−1^. Long-range electrostatic interactions were treated using the particle-mesh Ewald (PME) summation method [54] with a real space cutoff of 1.2 nm, Fourier spacing of 0.125 nm, and fourth-order interpolation. Short-range van der Waals interactions were computed within a cut-off distance of 1.0 nm for a99SB-disp [50] and 1.2 nm for C36m [51]. For the a99SB-disp force field, a long-range dispersion correction for energy and pressure was applied, while for C36m a force-switching function was used to avoid abrupt cut-off effects. Covalent bonds involving hydrogen atoms were constrained using the LINCS algorithm [55].

Single polyglutamine molecules were placed in an expanded conformation in a rhombic dodecahedron box with periodic boundary conditions. The system was then filled with water, after which sodium and chloride ions were inserted into the simulation box by replacing water molecules to reach an ionic strength of 150 mM. The systems were then energy minimized using a steepest-decent algorithm, followed by a brief equilibration of the solvent under NVT (100 ps) and NPT (100 ps). To completely randomize the initial conformation, a quick temperature annealing was performed in which the temperature was linearly increased to 1000 K and decreased to 300 K during a 20 ns NVT simulation. Subsequently, an equilibration run of 50 ns was performed. To reduce the computational cost of the simulation, the simulation volume was then reduced such that there is at least a separation of 2.0 nm between periodic images at all times during the simulation. The system was resolvated and equilibrated again following the procedure described above (skipping the temperature annealing step), after which a production simulation was carried out for 500 ns.

We performed four replicas of polyQ18, 23, 28, 38, and 48 using both all-atom force fields. For each polyQ length, the radius of gyration was measured across the aggregate simulation trajectory of the eight replicas (black line in Fig. 1f). Although the two force fields show variation in protein dynamics (Suppl. Figs. S2 and S3), the average radius of gyration of the polyQ chains across the four replicas per force field was comparable. We used the average radius of gyration of all eight replicas for each polyQ length to parametrize the hydrophobic strength (λ_*i*_) of the BB and SC beads in the 2BPA model.

#### Coarse-grained simulations

Following this, we conducted simulations of polyQ chains using the 2BPA-Q model, systematically testing various combinations of BB and SC hydrophobicities to identify the combination that best captures the scaling behavior with polyQ length. Coarse-grained molecular dynamics simulations were performed with the GROMACS [44] software package (version 2019.6), with a modified implementation of the 3OUT virtual site scheme to construct the HB beads as defined in Eqs. 5 and 6. Simulations are conducted at 300 K using a time step of 0.02 ps and inverse friction coefficient *γ*^−1^ = 2 ps for the Langevin dynamics integrator.

We systematically sampled hydrophobicity values of both BB and SC beads in increments of 0.01. Simulations were initiated from an extended conformation and equilibrated for 500 ns with hydrogen bonds turned off. Following the equilibration step, production runs were performed for 5000 ns with hydrogen bonds activated (*ε*_HB_ = 6.6 kJ*=*mol), and the resulting data were used to calculate the average radius of gyration. For each BB hydrophobicity value, we identified the corresponding SC hydrophobicity that minimized the error, defined as the mean squared difference from the all-atom measurements (Suppl. Fig. S4). Very good agreement with the all-atom radii of gryation (cf. Fig. 1f) was obtained for (λ_BB_, λ_SC_) = (0.64, 0.58) (Fig. 1f).

### Analysis of aggregation simulations

To characterize the aggregation behavior using our 2BPA-Q model, we applied three complementary analysis methods: clustering analysis, hydrogen bonding analysis, and amyloid zipper analysis.

#### Clustering analysis

Clustering analysis was conducted using the built-in *gmx clustsize* utility in GROMACS, using a cutoff distance of 0.6 nm to define a cluster. Physically, this cut-off implies that two molecules are considered to be part of the same cluster if any of their BB or SC beads are within 0.6 nm of each other. This method provides the number of clusters during the simulation, offering insights into the aggregation state, from dispersed monomers to large aggregates.

#### Hydrogen bonding analysis

Hydrogen bonding was analyzed using an in-house Python script based on MDAnalysis [56]. We calculated contacts between hydrogen bonding donor beads (HBH) and acceptor beads (HBO), using a cutoff distance of 0.12 nm. This threshold represents the range within which a hydrogen bond is considered formed, with an ideal hydrogen bond corresponding to a distance of 0 nm between HBH and HBO. Both intra- and intermolecular hydrogen bonds were included in the analysis. To normalize the data, the total number of hydrogen bonds was divided by the number of molecules in the simulation, yielding the average number of hydrogen bonds per molecule. Each residue can participate in up to two hydrogen bonds, one through the HBH bead and one through the HBO bead.

#### Amyloid zipper analysis

The steric zipper conformation was identified by analyzing residue-level contacts between BB and SC beads. Two criteria were used to classify residues as being in a steric zipper conformation:

1. The distance between the closest BB and SC beads must be between 0.38 nm and 0.5 nm.
2. The distance between the farthest BB and SC beads must be between 1.2 nm and 2.5 nm.

The first criterion ensures that the two residues are in close proximity, while the second criterion ensures that their side chains are aligned in the same direction, as expected in a steric zipper conformation. The cutoffs for these criteria, illustrated in Suppl. Fig. S5, were chosen by trial and error to ensure accurate identification of residues that are in the steric zipper conformation. To normalize the data, the total number of amyloid zipper residues was divided by the total number of residues in the simulation, yielding the percentage of residues in the amyloid state. The analysis was performed using the *contact matrix* pipeline from MDAnalysis [56], and the identified zipper residues were verified using VMD visualization software [57].

## Results

### Balance of specific and non-specific interactions determines aggregation behavior

To thoroughly assess the capabilities and versatility of our model beyond the calibrated parameter set, we conducted an extensive exploration by constructing a “phase diagram” that maps the effect of both side chain hydrophobicity (λ_SC_) and hydrogen bonding strength (*ε*_HB_). This approach involved generating 35 unique parameter combinations (λ_SC_, *ε*_HB_), arranged in a 5×7 grid. For each combination, we initiated simulations starting from a homogeneous solution of polyQ monomers, allowing us to observe how different parameter settings influence the aggregation behavior. This systematic variation enabled us to probe the model’s capacity to capture a wide range of aggregation pathways, providing insights into the underlying mechanisms of polyQ aggregation.

Our parameter sweep across the λ_SC_-*”*_HB_ space was centered on the default parameters of the 2BPA-Q model that were obtained from the calibration against the all-atom simulations. The sweep spanned values of λ_SC_ from 0.48 to 0.68 (in increments of 0.05) and values of *ε*_HB_ from 4.6 to 10.6 kJ*=*mol (in increments of 1.0 kJ*=*mol). In each simulation, 100 polyQ48 molecules were randomly distributed within the simulation box at a molar concentration of 1.0 mM. The simulations were run for 5.0 µs (2.5 × 10^8^ steps), at which point the systems that underwent aggregation were fully aggregated.

Reaching full equilibrium, where all molecules form a single cluster, likely requires significantly longer simulation times. Our simulations already provide a qualitative picture of how the systems evolve under different parameter conditions.

Our results reveal that the balance between specific (hydrogen bonding) and nonspecific (hydrophobic) interactions plays a critical role in determining polyQ aggregation behavior. By systematically varying λ_SC_ and *ε*_HB_, we observed distinct aggregation pathways depending on how these interactions were balanced. When nonspecific hydrophobic interactions dominate (λ_SC_ ≥ 0.63), phase separation occurred consistently, resulting in the formation of clusters without any free monomers, irrespective of the hydrogen bonding strength (Fig. 2b). This indicates that side chain hydrophobicity alone is sufficient to drive clustering, leading to the condensed, nonspecific aggregation of polyQ molecules. However, without sufficiently strong hydrogen bonding, these clusters lacked the ordered structure characteristic of amyloid fibrils.

**Fig. 2.**
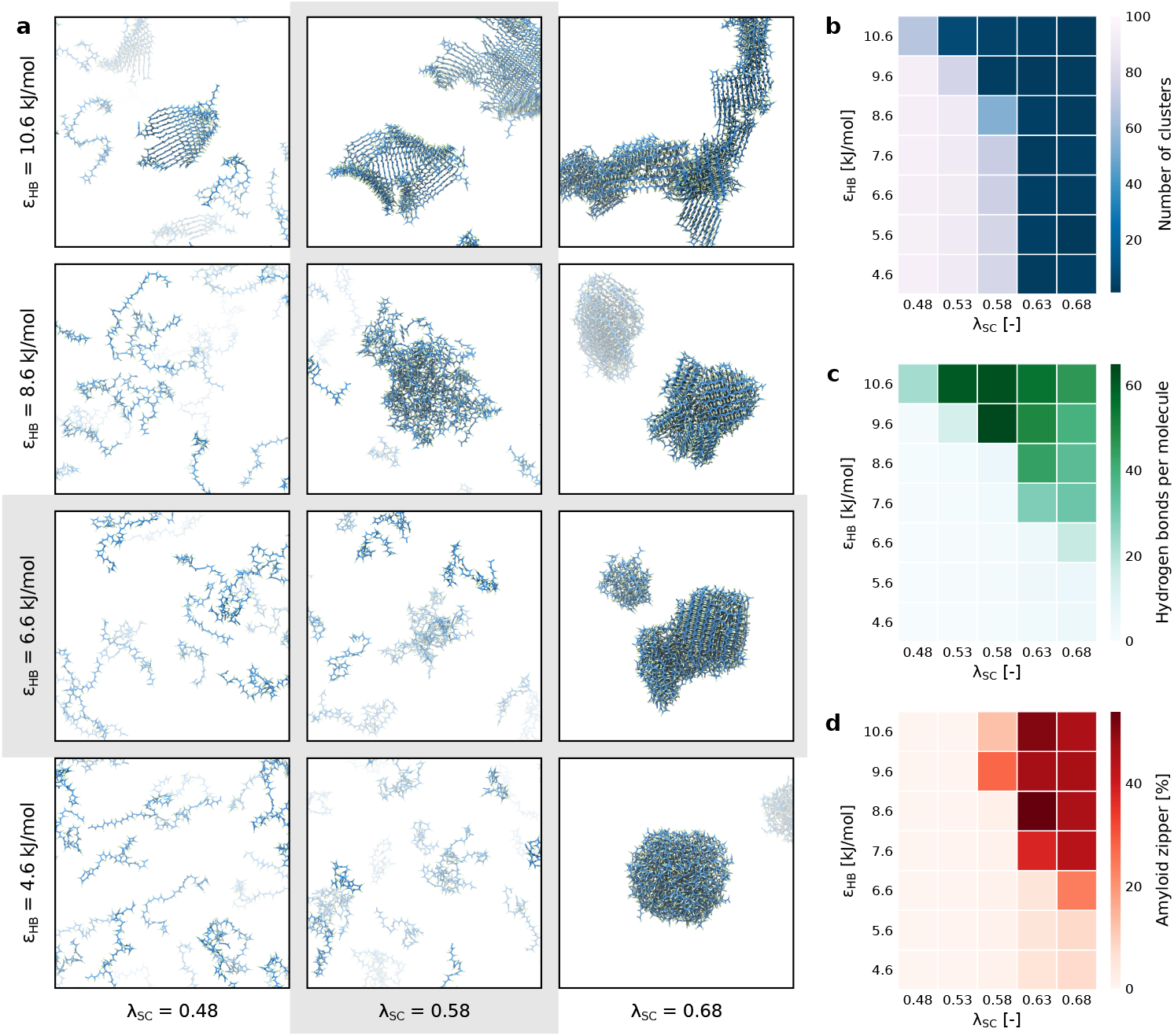
PolyQ48 aggregation behavior for various combinations of side chain hydrophobicity λ_SC_ and hydrogen bonding strength *ε*_HB_. **Phase diagram of Q48 aggregation behavior as a function of side chain hydrophobicity (**λ_SC_**) and hydrogen bonding strength (***ε*_HB_**) on polyQ aggregation behavior**. (**a**) Representative snapshots of polyQ structures at *t*^∗^ = 5 µs at various parameter combinations (λ_SC_; *ε*_HB_). The phase diagram displays diverse aggregation behaviors, including no aggregation (bottom left), spherical condensates (bottom right), β-sheets (top left), and amyloid fibers (top right). (**b**) Clustering analysis of each simulation, with clustering values representing the number of clusters. (**c**) Hydrogen bonding analysis of each simulation, with hydrogen bonding values representing the average number of hydrogen bonds per molecule. Note that each residue can participate in maximum two hydrogen bonds, one through the HBH bead and one through the HBO bead. (**d**) Amyloid zipper analysis of each simulation, with amyloid zipper values representing the fraction of residues that is in a steric zipper conformation.

In contrast, increasing the hydrogen bonding energy facilitated the formation of more ordered structures, such as β-sheets and amyloid fibers. For *ε*_HB_ ≥ 7.6 kJ*=*mol and λ_SC_ ≥ 0.63, the systems consistently aggregated into amyloid fibers (Fig. 2d). However, the aggregation pathway varied: some systems aggregated via hydrophobic interactions and phase separation, while others at λ_SC_ = 0.58 first formed large β-sheets that later transitioned into amyloid fibers (Suppl. Fig. S7 and S8). This highlights that while specific interactions through hydrogen bonding are essential for stabilizing amyloid structures, they require hydrophobic interactions to concentrate polyQ molecules prior to amyloid formation.

The balance between these interactions becomes most apparent in extreme cases of either hydrogen bonding energy or side chain hydrophobicity. For example, at *ε*_HB_ = 10.6 kJ*=*mol and λ_SC_ = 0.53, large β-sheets and β-barrels formed but did not progress to amyloid zippers due to insufficient hydrophobic interactions. Conversely, at *ε*_HB_ = 6.6 kJ*=*mol and λ_SC_ = 0.68, polyQ molecules initially phase separated into condensates, which gradually matured into amyloid fibers (Suppl. Fig. S8). This demonstrates that aggregation pathways are controlled by a delicate balance: hydrophobic clustering brings molecules together, while specific hydrogen bonding induces and stabilizes the amyloid structure.

Interestingly, our simulation using the default parameters (obtained from the calibration process) reveals that the system appears to be near a critical point in the phase diagram, where it remains disordered but is close to both aggregation pathways: phase separation driven by side chain hydrophobicity and β-sheet formation through hydrogen bonding. Although no strong phase separation occurs, we observed the formation of small transient clusters rather than large stable aggregates. Similarly, although amyloids are stable at the default hydrogen bonding energy (Fig. 1e), we do not observe the formation of stable β-sheets when starting from a solution of polyQ monomers. This indicates that the system is near a tipping point between different aggregation states, where the interactions are sufficient to promote some degree of clustering but not strong enough to drive the formation of ordered amyloid structures. Indeed, the disorder-to-order transition of polyQ monomers is a rare event [33, 58], requiring extensive sampling to capture in simulations. The results obtained from the phase diagram highlight the sensitivity of the system to slight changes in the balance of specific and nonspecific interactions, placing the default parameters near the boundary between disorder and structured aggregation.

### Two modes of amyloid growth

During seeded aggregation, polyQ monomers and small oligomers dock onto an amyloid segment and undergo a disorder-to-order conformational transition, adopting the amyloid structure. Understanding the mechanisms by which monomers attach to existing amyloid segments is crucial to unraveling the aggregation process. To explore these growth mechanisms, we conducted controlled seeded aggregation simulations, introducing a pre-built polyQ amyloid (a 4×4 antiparallel stack of Q23; see Suppl. Fig. S9a) into a simulation box containing Q23 monomers. Our simulations revealed two distinct amyloid growth mechanisms: β-sheet elongation, where polyQ monomers form hydrogen bonds with existing β-sheets, thereby extending the β-sheets, and the “zipper mechanism” [58], where the side chains of the monomers interlock with those of the β-sheets, adding a new β-sheet to the amyloid structure.

Focusing first on β-sheet elongation, this process begins with a polyQ monomer approaching an amyloid segment, drawn in by hydrophobic interactions. Initially, the monomer docks in a disordered state (Fig. 3a). Through random fluctuations, it starts to form hydrogen bonds with the amyloid. However, individual hydrogen bonds are too weak to stabilize the connection, causing the bonds to form and break repeatedly. Eventually, the monomer aligns so that several consecutive residues form stable hydrogen bonds with one of the β-sheets of the amyloid (Fig. 3b). This stabilizes the interaction, allowing more hydrogen bonds to form, and the monomer becomes a strand of the β-sheet. Over time, the monomer stabilizes further by forming more hydrogen bonds, thus becoming fully integrated in the amyloid segment (Fig. 3c).

**Fig. 3.**
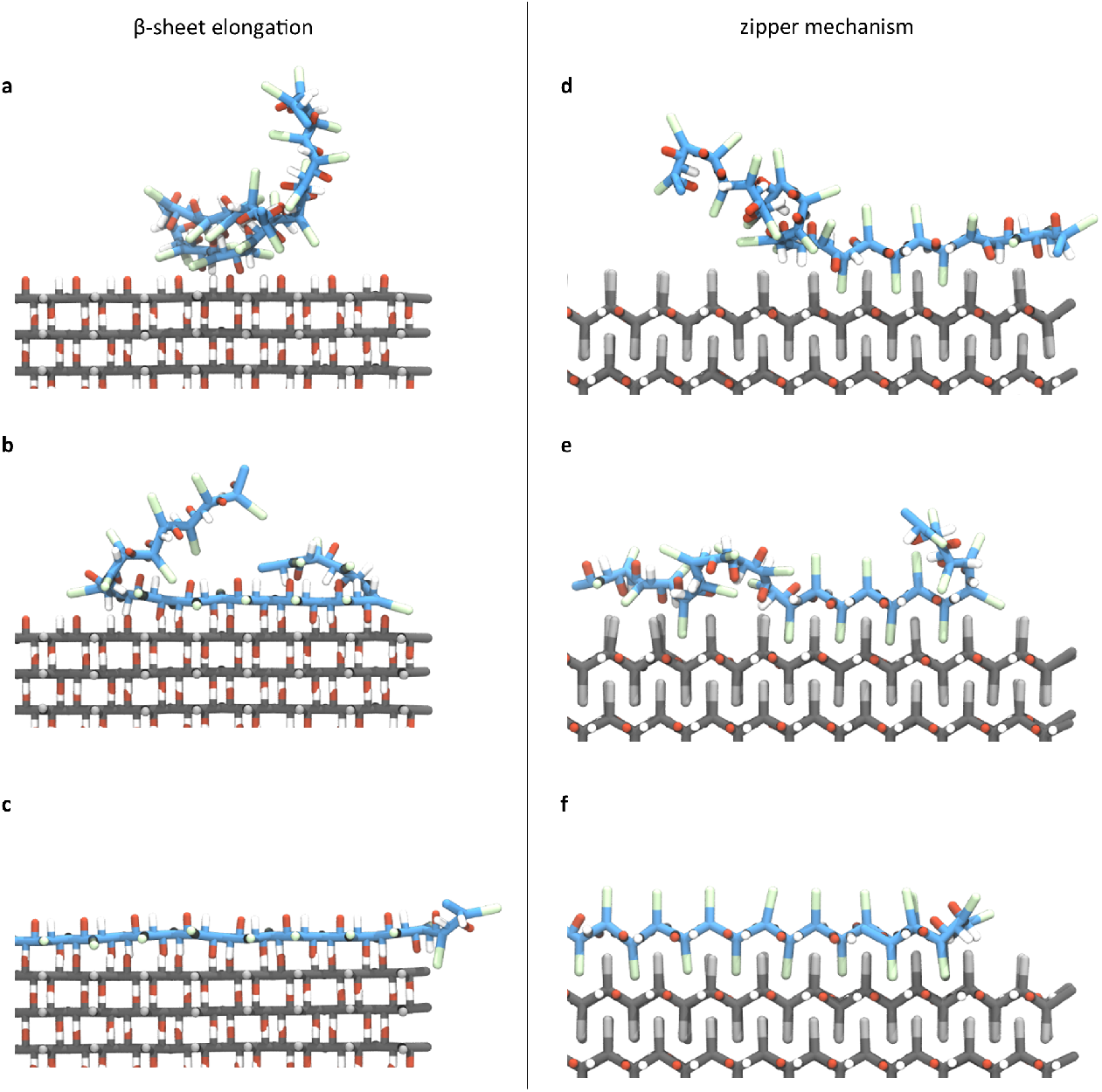
Two modes of amyloid growth. (**a**–**c**) Amyloid growth through β-sheet elongation. A polyQ monomer (Q23, blue) attaches to an amyloid segment (gray) by forming hydrogen bonds with one of the β-sheets of the amyloid. (**d**–**f**) Amyloid growth through the zipper mechanism. A polyQ monomer (blue) attaches to an amyloid segment (gray) by interlocking its side chains with the side chains of the amyloid. Trajectory animations of both growth modes are available in Suppl. Movies S1 and S2.

Distinct from β-sheet elongation, amyloid segments also grow by adding new β-sheets through the zipper mechanism. As a monomer approaches the amyloid, it docks via hydrophobic interactions. Through fluctuations and hydrophobic interactions, one of the monomer’s side chains slips between two consecutive side chains of the amyloid. Initially, this connection is unstable, leading to repeated dissociation and reassociation of the monomer (Fig. 3d). However, when the monomer side chains become kinetically trapped between the side chains of an existing β-sheet, the interaction becomes more stable, initiating a locking mechanism (Fig. 3e). As more side chains are interlocked in a zipper-like fashion, the monomer is firmly attached to the amyloid (Fig. 3f). This connection is highly stable, making dissociation unlikely.

### Seeded aggregation simulations reveal distinct growth kinetics for Q48 and Q23 monomers

To further explore the mechanism of polyQ aggregation, we performed additional seeded aggregation simulations in which we placed a small amyloid core (comprising a 4×4 array of Q23 β-strands) in a simulation box with 200 disordered polyQ monomers at a molar concentration of 1.0 mM (Suppl. Fig. S9b). We conducted simulations for both Q48 and Q23 monomers under identical conditions. To maintain a consistent concentration in the dilute phase throughout the simulation, we paused the simulation every 100 ns to measure the size of the largest cluster (i.e., the preformed seed plus any attached monomers) using the gmx clustsize utility. If the concentration of the dilute phase decreased, we restored it back to 1.0 mM by adding additional monomers. Each simulation was run for a total of 10 µs, with five independent replicas for both the Q48 and Q23 systems to ensure reproducibility of the results.

In all simulation replicas, we observed substantial growth of the amyloid seed, with most of the growth occurring in the β-sheet elongation direction (Fig. 4a). Growth along the steric zipper direction, while present, was comparatively slower. For each system, the size of the largest cluster was tracked over time and the growth trajectories from the five replicas were averaged into a single growth curve for both Q48 and Q23 monomers (Fig. 4b and Suppl. Fig. S10). The results show a clear distinction in the aggregation kinetics between the two monomer lengths. Despite maintaining identical starting conditions and monomer concentrations, the growth of Q48 was significantly faster than that of Q23.

**Fig. 4.**
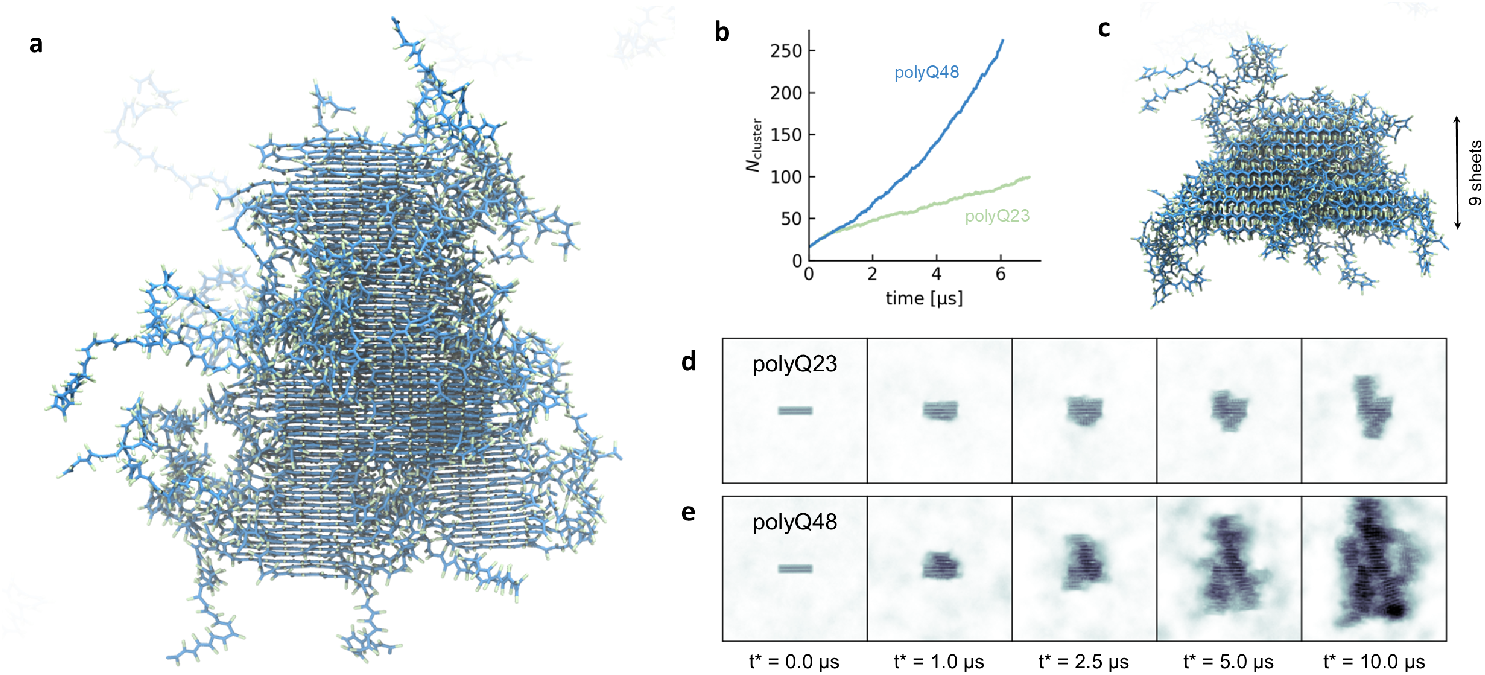
Diverse aggregation behavior for various combinations of side chain hydrophobicity λ_SC_ and hydrogen bonding strength *ε*_HB_. (**a**) PolyQ48 amyloid fiber after 5 µs of simulation time. The fiber has primarily grown in the β-sheet elongation direction. (**b**) Number of molecules that are part of the amyloid as a function time, for polyQ48 and Q23. The curves are averaged over five simulation replicas (Suppl. Fig. S10). (**c**) After 5 µs, the fiber has a thickness of nine β-sheets. (**d**) Evolution of the amyloid seed over time. Snapshots are obtained by aligning the simulation trajectory to the initial amyloid seed and averaging the density distribution over 0.1 µs. Density snapshots for all replicas are shown in Suppl. Figs. S11 and S12. An animation of the Q48 seeded simulation is available in Suppl. Movie S3.

Interestingly, fiber growth was symmetric along the β-sheet elongation axis, with both ends of the seed growing at similar rates. While the fibers elongated, we also observed an increase in thickness, which corresponds to the stacking of additional β-sheets perpendicular to the elongation direction (zippering). Starting with a seed composed of four β-sheets, most fibers reached a thickness of approximately nine β-sheets by the end of the 10 µs simulations (Fig. 4c). This consistent thickening highlights the ability of polyQ monomers to stack on top of the growing fiber core, contributing to the formation of well-ordered, multilayered amyloid structures and matches the structural model proposed by experiments [15].

## Discussion

In this study, we developed a CGMD model for polyQ aggregation. The model was first parameterized using a combination of atomistic simulation data and experimental observations. After establishing these parameters, we explored the broader potential of the model by constructing a phase diagram, systematically varying hydrophobicity and hydrogen bonding strength beyond the calibrated parameter values. This allowed us to map out the range of possible aggregation pathways the model can display, highlighting its ability to capture both phase separation and amyloid formation. This approach provided new insights into the interplay between specific (hydrogen bonding) and nonspecific (hydrophobic) interactions that govern polyQ aggregation, revealing how small changes in interaction strengths can shift aggregation kinetics and pathways.

In the phase diagram simulations, we found that subtle increases in side chain hydrophobicity (λ_SC_) drive phase separation, which, depending on the strength of the hydrogen bonds (*ε*_HB_), can lead to the transition from a liquid to a solid-like aggregate state. Specifically, at sufficiently high hydrogen bonding strengths, the hydrophobic clustering of polyQ molecules transitions into ordered aggregates, including large β-sheets and amyloid fibrils. However, with the calibrated parameters obtained from atomistic simulations, the system did not undergo phase separation, nor did it spontaneously form amyloid fibers within the simulation time frame (up to 25 µs; Suppl. Fig. S13). This suggests that while the calibrated model accurately captures the critical conditions required for aggregation, the nucleation process, which typically initiates amyloid formation, is slow with a lag phase that is longer than the simulation time. Notably, while spontaneous nucleation was not observed, a preformed amyloid seed remained stable, and when placed in a solution of disordered polyQ monomers, it successfully induced seeded aggregation.

Our seeded aggregation simulations demonstrated that aggregation can occur by two mechanisms: β-sheet elongation and steric zippering, the former being the most dominant in our simulations. Importantly, we found that Q48 aggregates much faster than Q23, despite being held at the same molar concentration. This suggests that the length of the polyQ chain is a critical factor influencing the aggregation kinetics, with longer chains having a higher propensity for integration into growing amyloid fibers. Furthermore, at the relatively high concentrations used in our simulations, we observed branching of the fibers, indicating that concentration not only accelerates growth, but may also contribute to secondary nucleation [59].

Recent experimental work has demonstrated that the polyQ core is the primary driver of seeded aggregation in huntingtin (htt), even in the presence of flanking domains [60]. This finding underscores the relevance of our polyQ-centric model, which focuses exclusively on glutamine residues, to capture critical aspects of seeding behavior and aggregation. While not essential, the flanking domains of htt are known to modulate aggregation behavior [11, 15]. Our model currently omits these domains, but it can potentially be extended to incorporate htt-specific features, further enhancing its predictive power. Such extensions would allow for a more comprehensive exploration of htt aggregation mechanisms in future studies.

Despite its simplicity, our coarse-grained model has proven to be a powerful tool for simulating polyQ monomer aggregation into amyloid fibers. Future improvements could focus on refining the side chain representation, replacing the current simplistic approach with a more detailed model that allows for multiple side chain conformations. This would enable better representation of disordered monomers and the formation of amyloid fibers [41]. Furthermore, the model could be expanded to include all 20 amino acids, each with distinct side chain properties such as length and hydrophobicity, making it applicable to a wide range of aggregation-related studies beyond polyQ.

In conclusion, our study presents a coarse-grained molecular dynamics model that successfully captures key aspects of polyQ aggregation, including phase separation, β-sheet formation, and seeded amyloid growth. Through systematic parameter exploration and seeded aggregation simulations, we demonstrated that the balance between specific and nonspecific interactions is crucial to determine the aggregation behavior. The model accurately predicts many features of polyQ aggregation, and its simplicity offers a powerful basis for refinement and extension, offering a promising platform for future studies on protein aggregation and amyloid-related diseases.

## Acknowledgements

This work was financially supported by the Netherlands Organization of Scientific Research grant no. OCENW.GROOT.2019.068. The authors thank Vasista Adupa for assistance with all-atom molecular dynamics simulations. The authors thank the Center for Information Technology of the University of Groningen for their support and for providing access to the Hábrók high-performance computing cluster.

---

**Suppl. Fig. S1.**
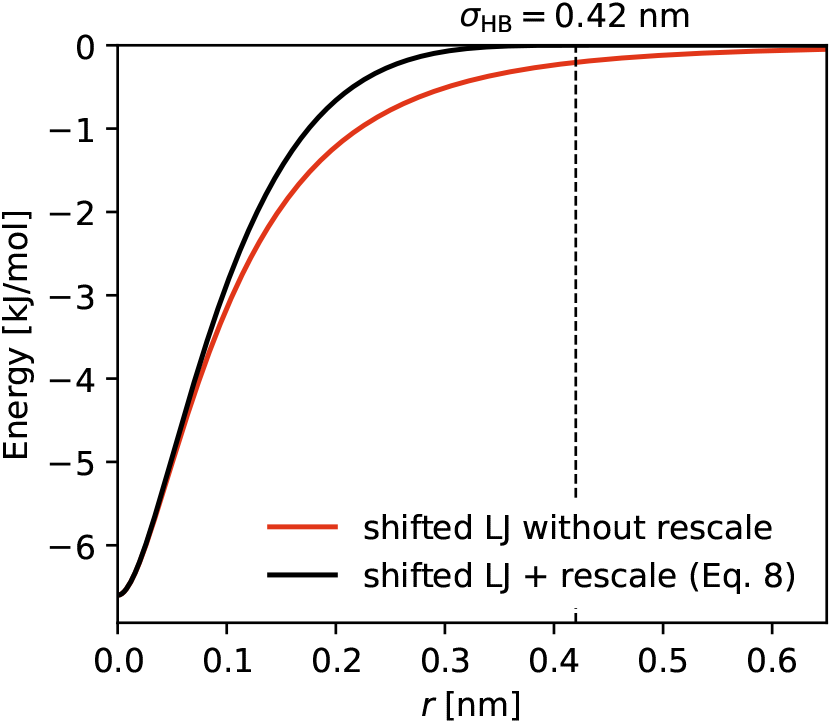
Effect of the scale function on the hydrogen bonding potential. The scale function (Eq. 9) ensures that the potential is zero at *σ*_HB_ = 0.42 nm.

**Suppl. Fig. S2.**
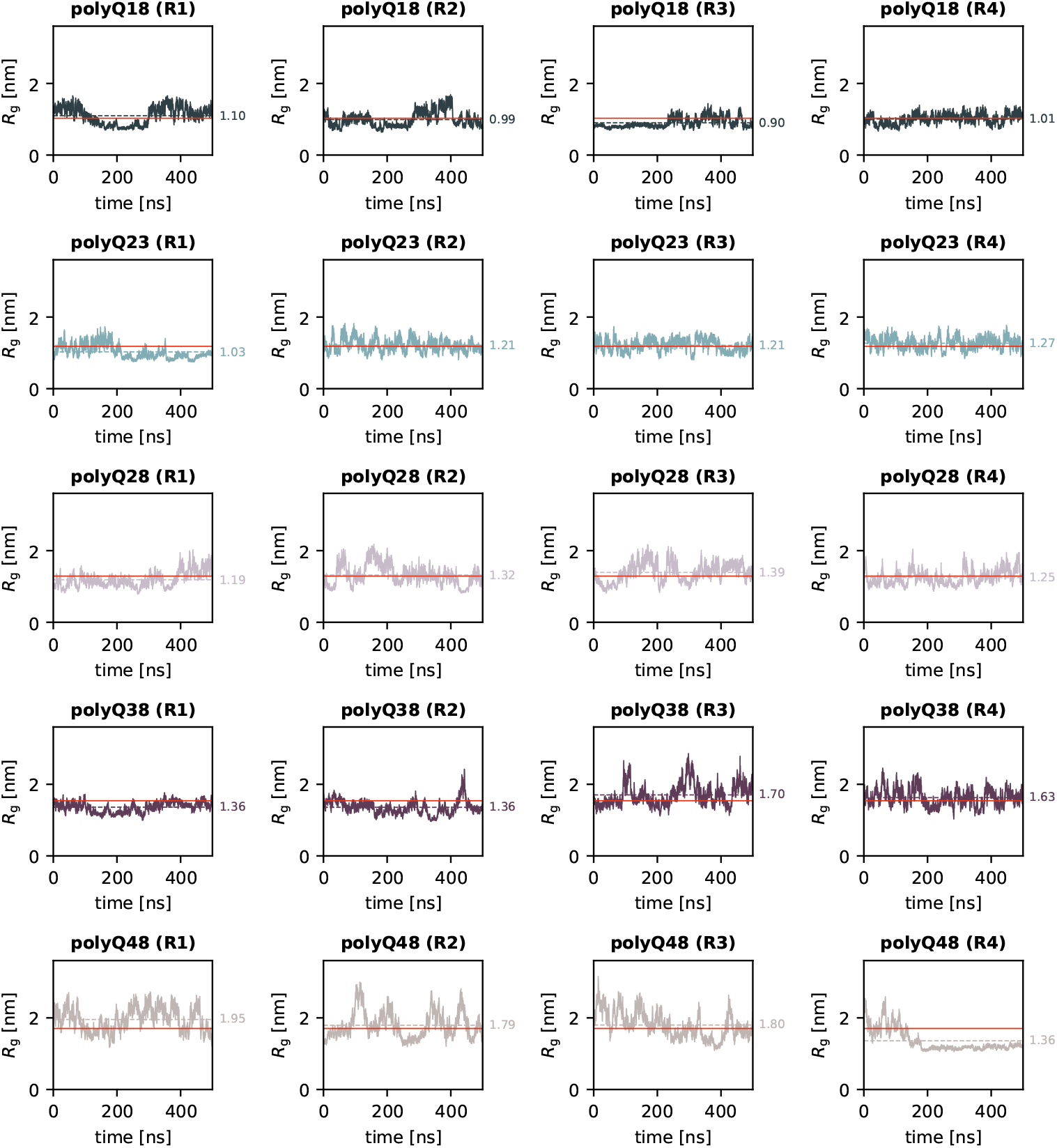
Radius of gyration of monomeric polyglutamine (amber99SB-disp). The dashed lines indicate the average radius of gyration of each replica simulation, the solid red line marks the average across all eight replicas (both a99SB-disp and C36m) for that particular polyQ length; see black markers in Fig. 1f.

**Suppl. Fig. S3.**
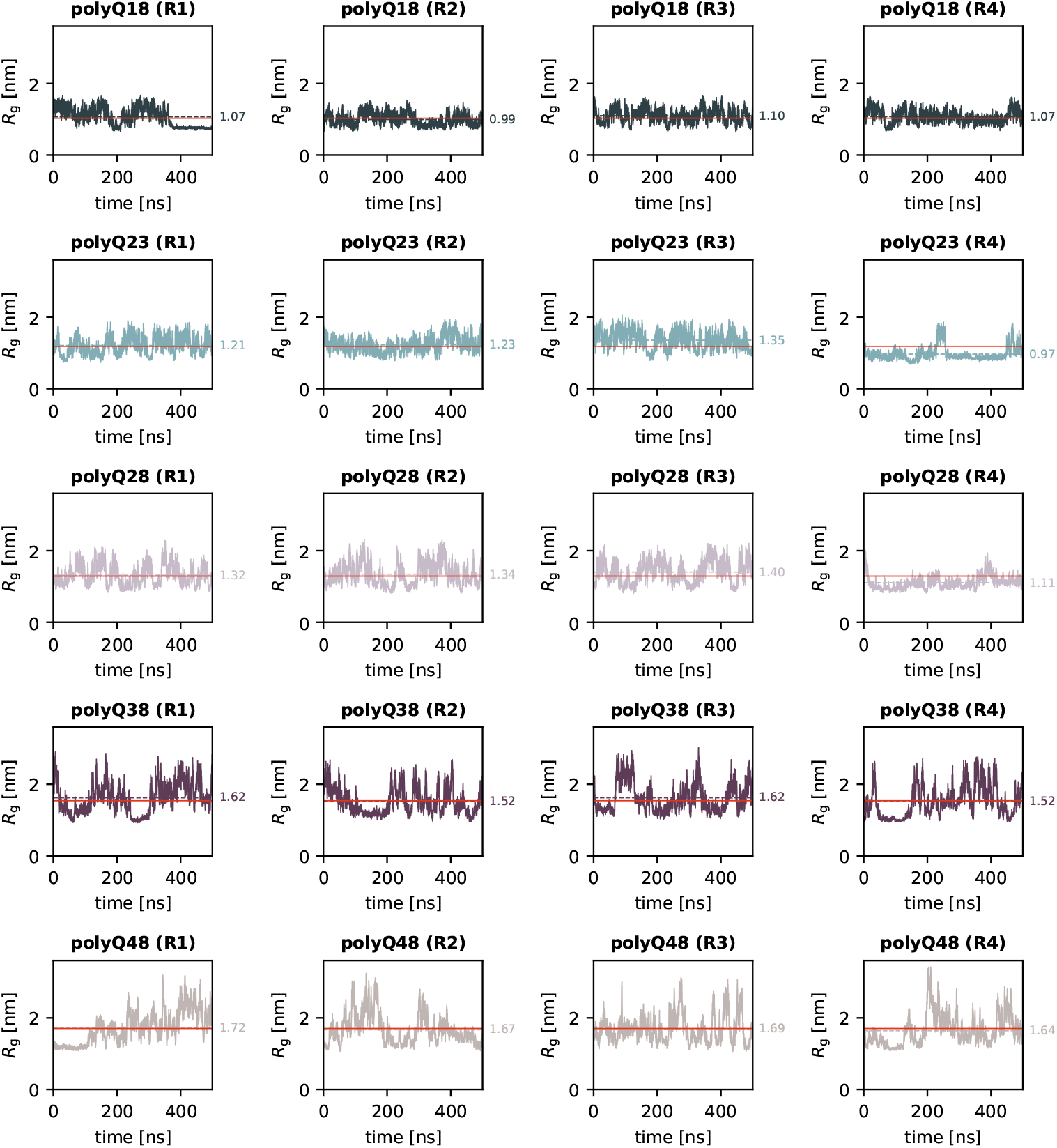
Radius of gyration of monomeric polyglutamine (CHARMM36m). The dashed lines indicate the average radius of gyration of each replica simulation, the solid red line marks the average across all eight replicas (both a99SB-disp and C36m) for that particular polyQ length; see black markers in Fig. 1f.

**Suppl. Fig. S4.**
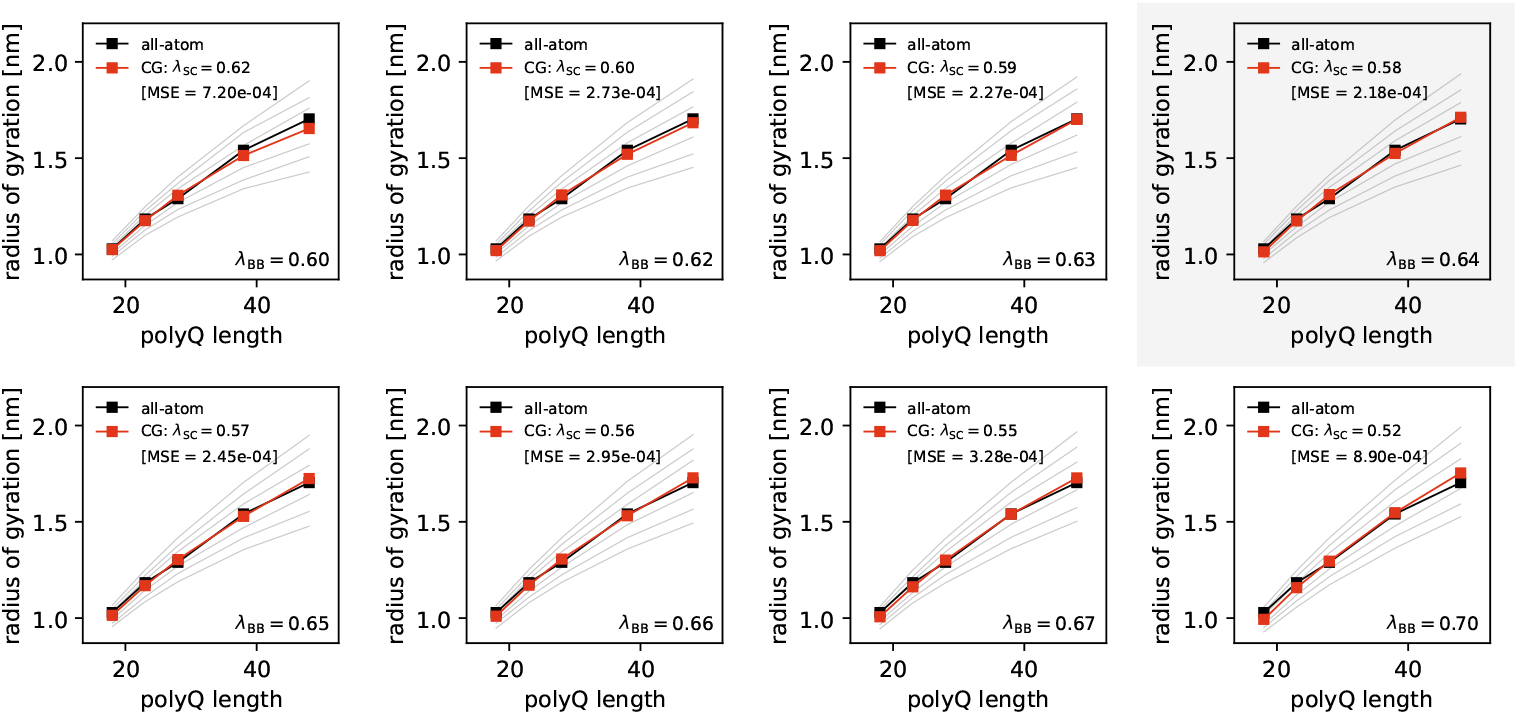
Systematic calibration of backbone and side chain hydrophobicity in the 2BPA-Q model. For each backbone hydrophobicity λ_BB_ the *R*_g_ scaling is calculated for different side chain hydrophobicities (λ_SC_ increments of 0.01, gray lines) and the optimal side chain hydrophobicity is determined (red lines) by calculating the mean squared error from the all-atom measurements. There are multiple combinations of λ_BB_ and λ_SC_ that very accurately capture the *R*_g_ length dependence of polyQ. The optimal combination of (λ_BB_, λ_SC_) is where the MSE is the smallest, i.e. (λ_BB_, λ_SC_) = (0.64, 0.58).

**Suppl. Fig. S5.**
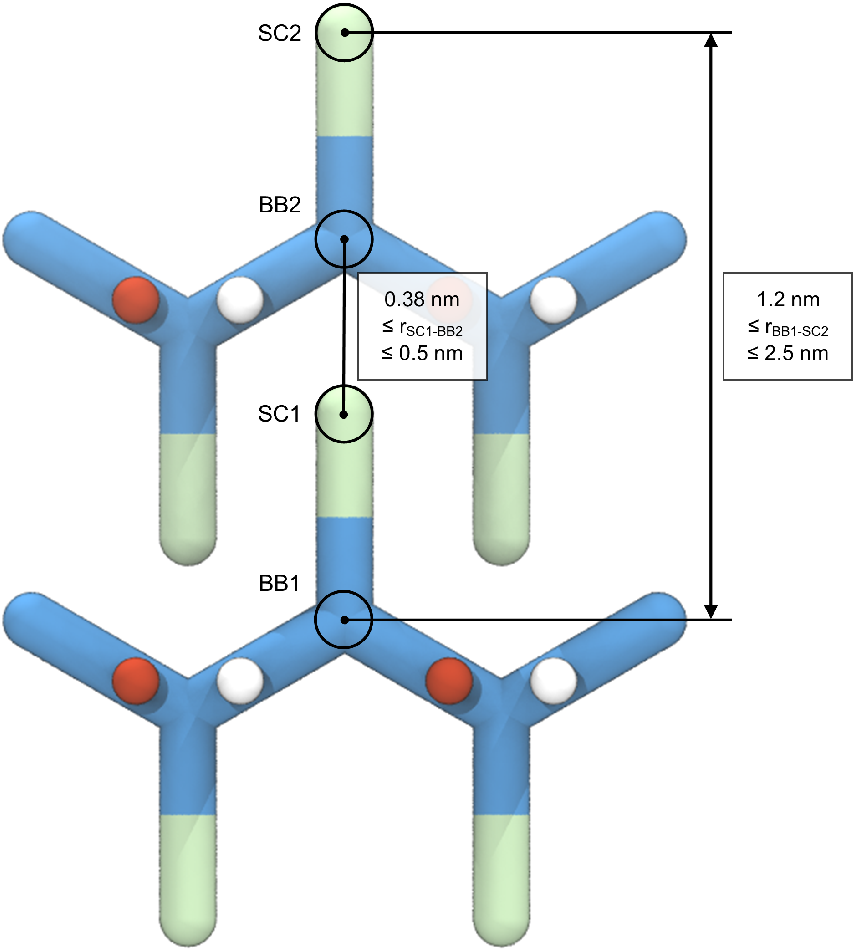
Distance criteria for zipper conformation. Two distance conditions have to be satisfied for two residues to be in steric zipper conformation.

**Suppl. Fig. S6.**
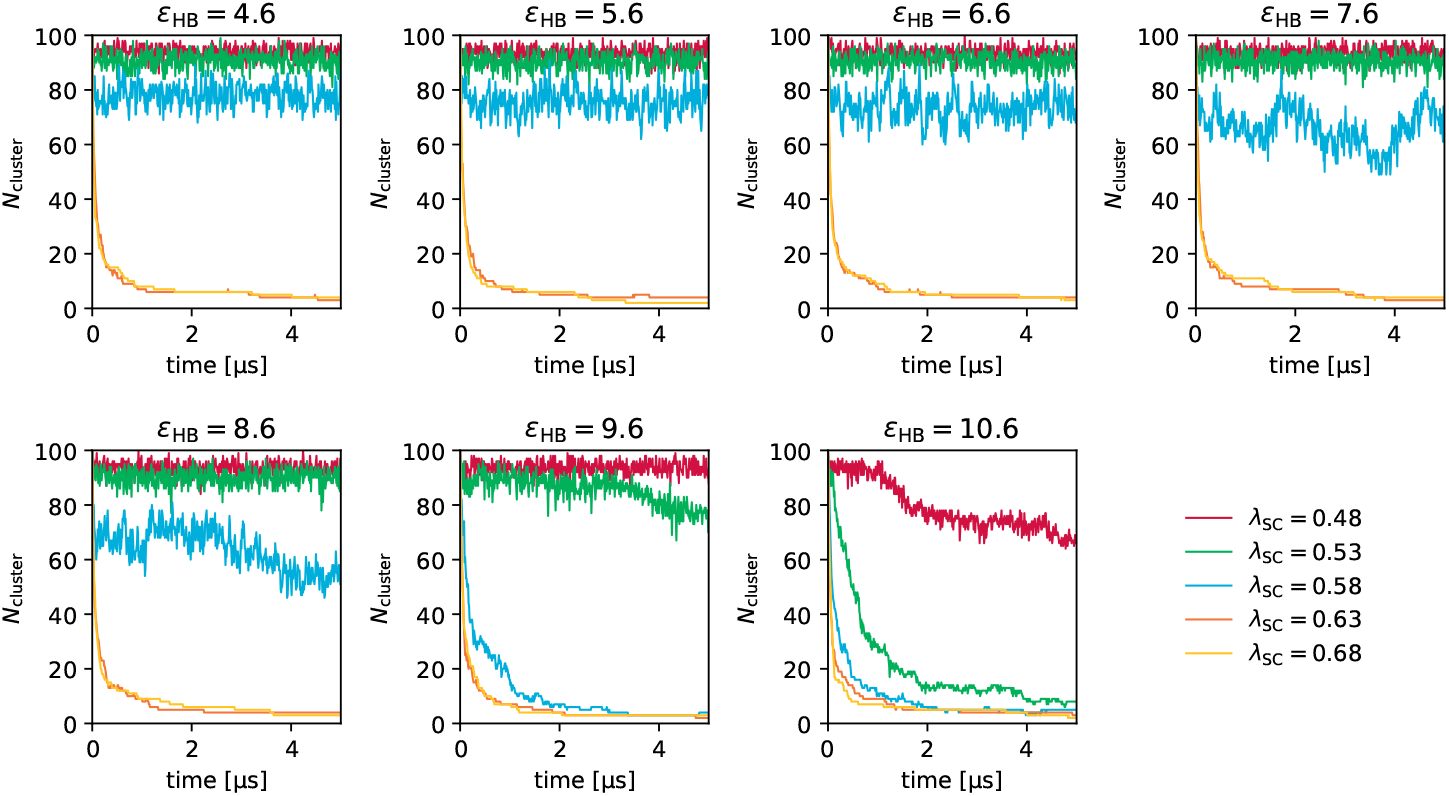
Number of clusters as a function of time for all combinations of (λ_BB_, λ_SC_) in the phase diagram. For *ε*_HB_ ≤ 8.6 kJ*=*mol, the clustering behavior is primarily dictated by side chain hydrophobicity (λ_SC_), with minimal influence from hydrogen bonding. At stronger hydrogen bonding energies (*ε*_HB_ = 9.6 and 10.6 kJ*=*mol), the λ_SC_ threshold for cluster formation decreases, indicating a synergistic effect between hydrogen bonding and hydrophobic interactions in driving cluster formation.

**Suppl. Fig. S7.**
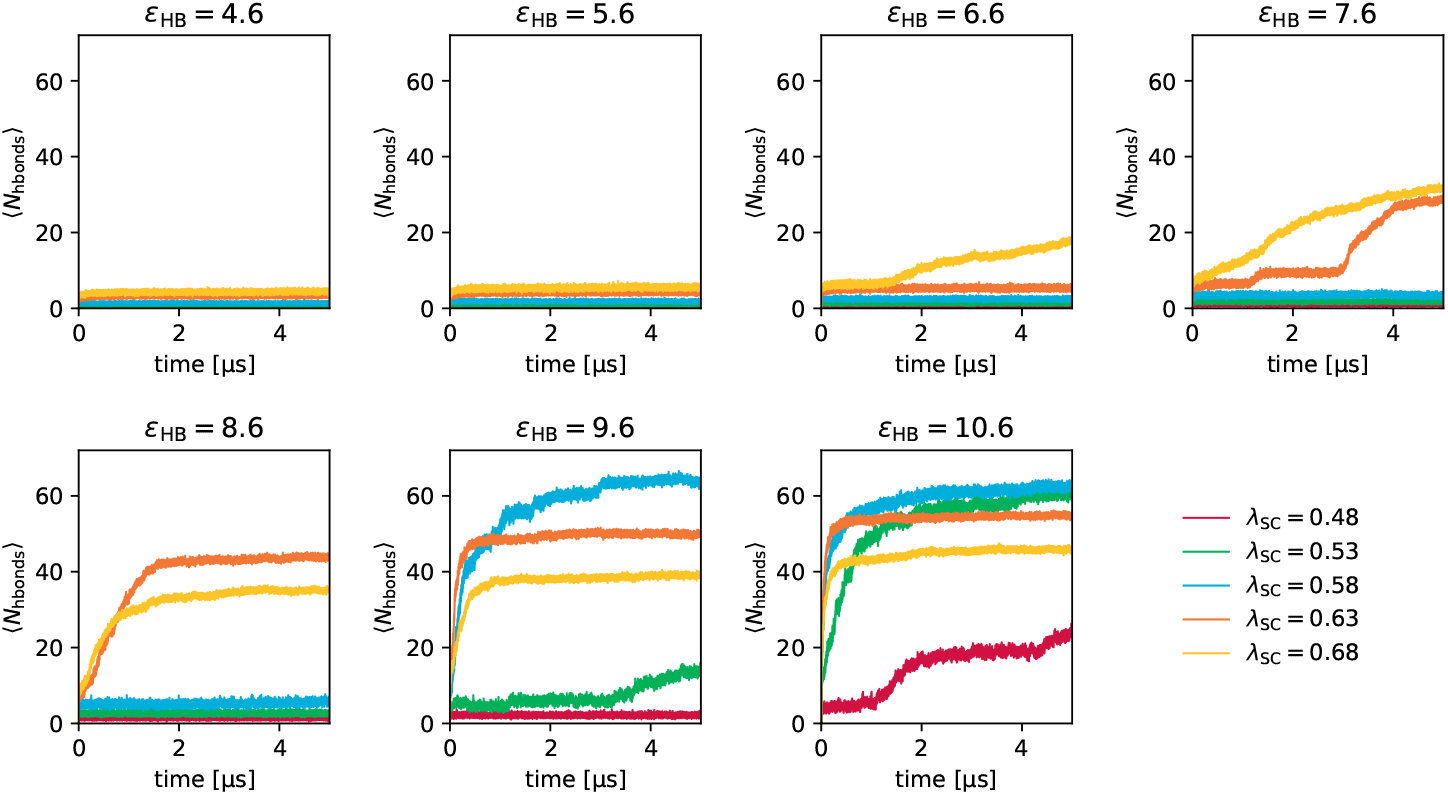
Average number of hydrogen bonds per molecule as a function of time for all combinations of (λ_BB_, λ_SC_) in the phase diagram. At low hydrogen bonding energies (*ε*_HB_ ≤ 5.6 kJ*=*mol), no hydrogen bonds are formed. As *ε*_HB_ increases, hydrogen bonds form only after molecules have clustered due to side chain hydrophobicity. At stronger hydrogen bonding energies (*ε*_HB_ = 9.6 and 10.6 kJ*=*mol), different aggregation pathways are observed. For hydrophobic side chains (λ_SC_ ≥ 0.63), clusters primarily form due to side chain hydrophobicity, while at lower hydrophobicities, both hydrogen bonding and hydrophobic interactions contribute to cluster formation, resulting in distinct aggregation behaviors and hydrogen bonding evolution curves.

**Suppl. Fig. S8.**
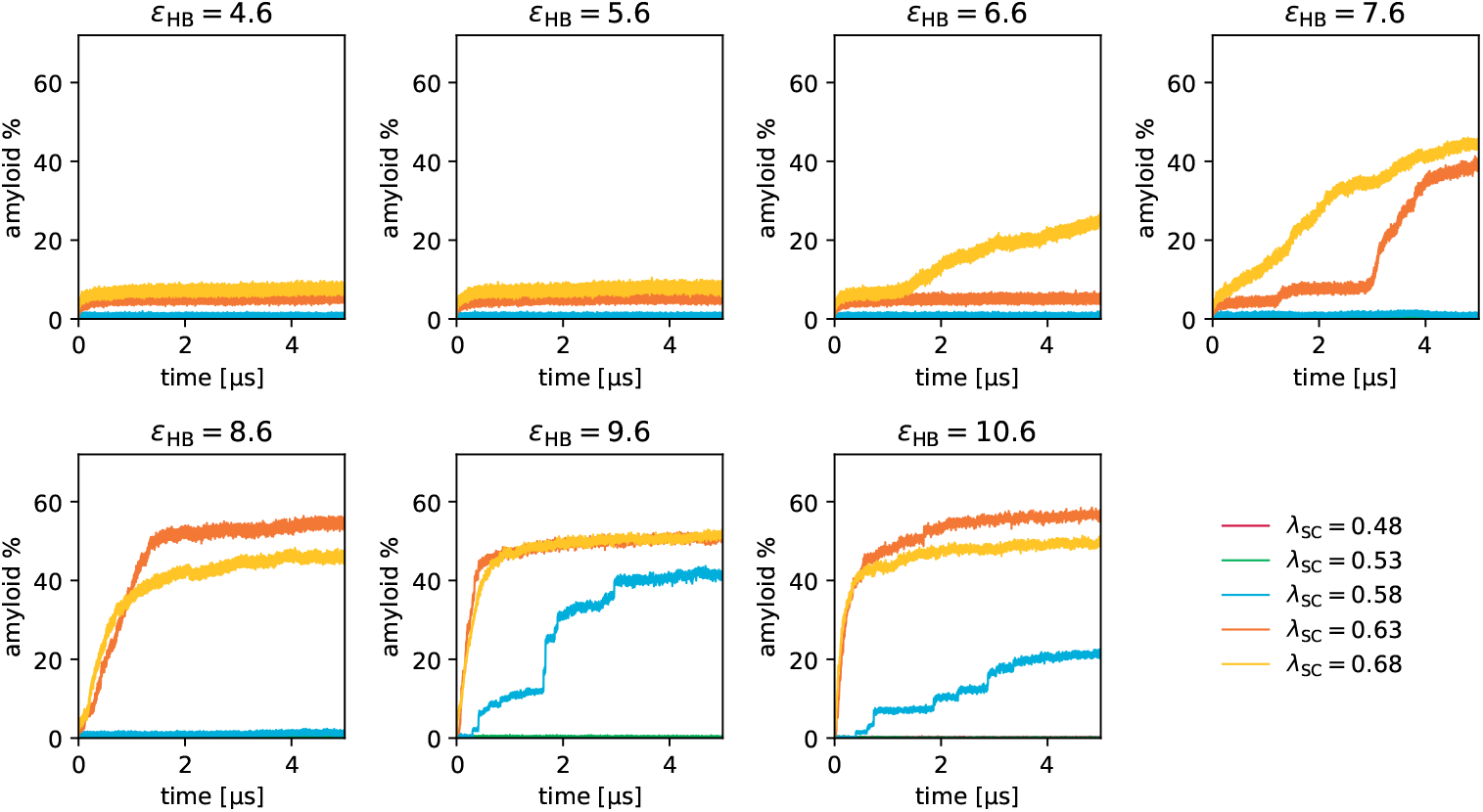
Amyloid formation as a function of time for all combinations of (λ_BB_, λ_SC_) in the phase diagram. At low hydrogen bonding energies (*ε*_HB_ ≤ 5.6 kJ*=*mol), no steric zippers are formed. As *ε*_HB_ increases, steric zippers form only after molecules have clustered due to side chain hydrophobicity, resulting in aggregation curves similar to those observed in the hydrogen bonding analysis. At stronger hydrogen bonding energies (*ε*_HB_ = 9.6 and 10.6 kJ*=*mol), steric zippers form only when the side chain hydrophobicity is sufficiently high (λ_SC_ ≥ 0.58). For lower λ_SC_ values, β-sheets form, but steric zippers do not.

**Suppl. Fig. S9.**
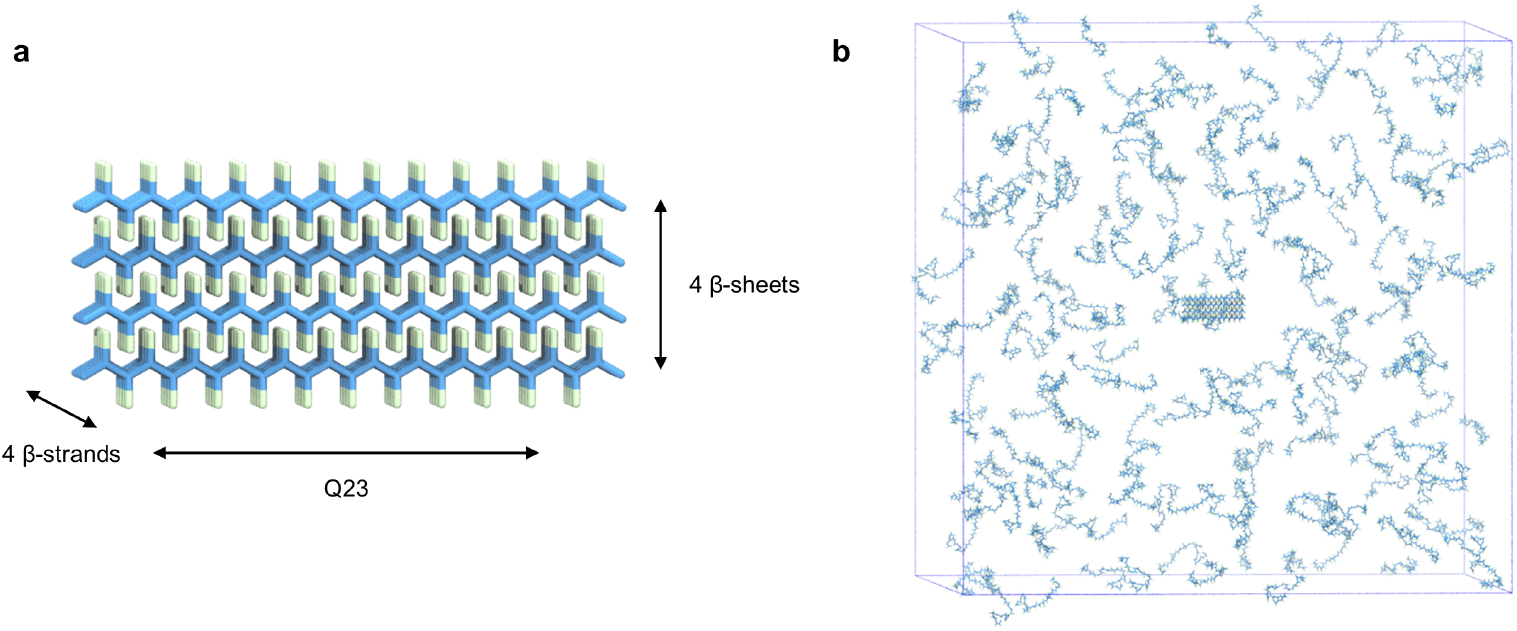
Setup of seeded simulations. (**a**) The amyloid core used in the seeded simulations consists of a 4×4 antiparallel stack of Q23 monomers; hydrogen bonding beads are omitted for clarity. (**b**) The seed is placed in a simulation box containing 200 randomly distributed polyQ monomers at a molar concentration of 1.0 mM (box dimensions: 70 nm). The setup shown here corresponds to Q48 seeded simulations. Note that the amyloid core remains the same (4×4 antiparallel stack of Q23).

**Suppl. Fig. S10.**
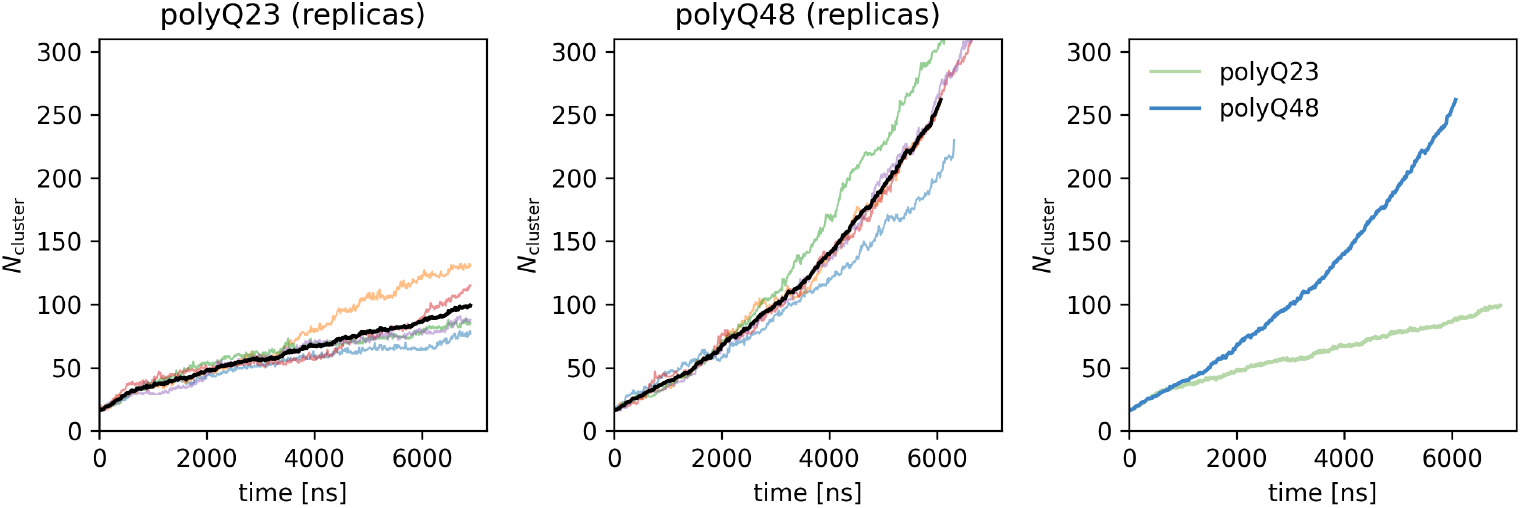
Size of the largest cluster (number of molecules) for each of the replicas in the seeded aggregation simulations.

**Suppl. Fig. S11.**
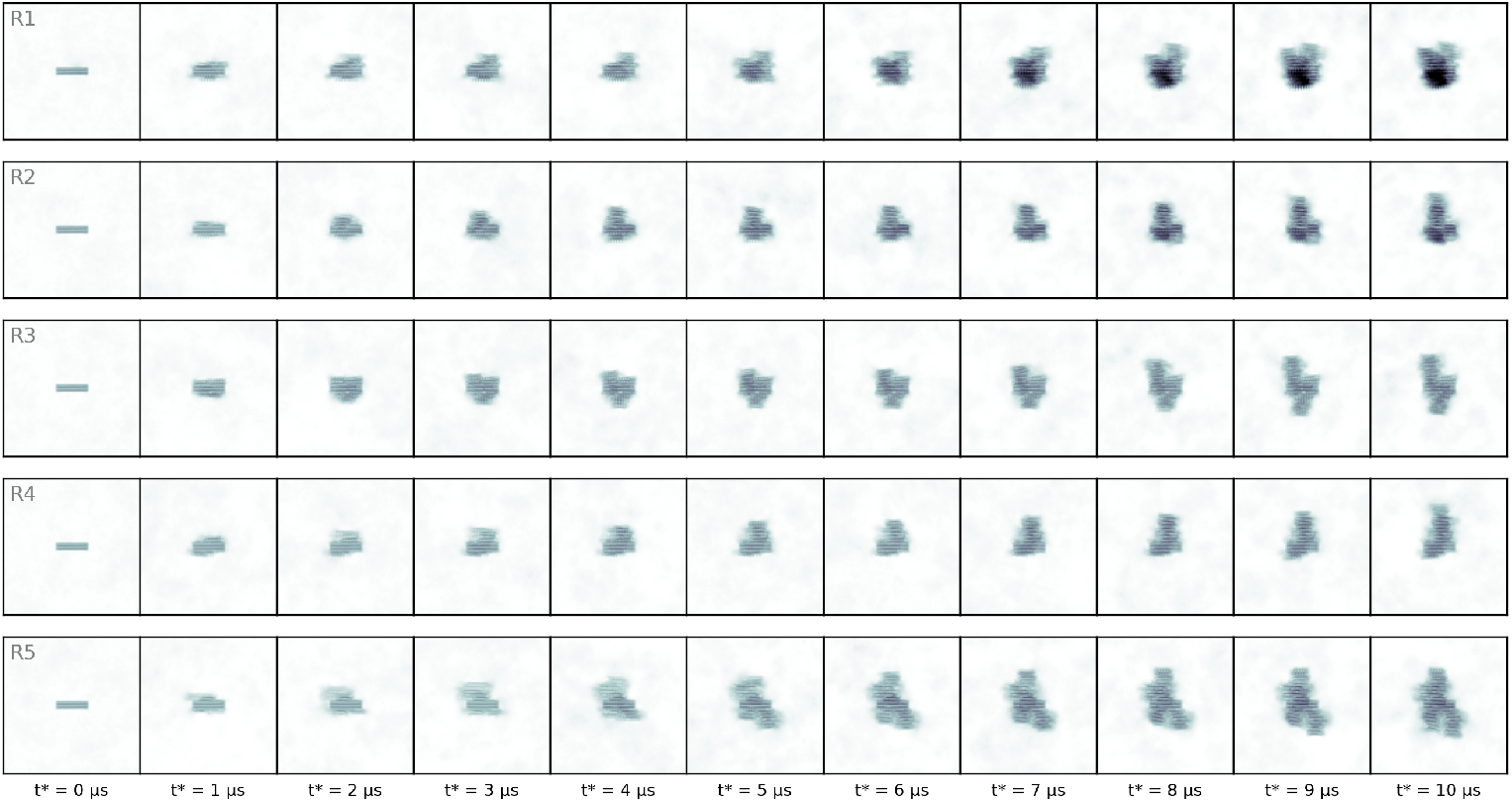
Amyloid growth for five replicas of seeded simulations (polyQ23).

**Suppl. Fig. S12.**
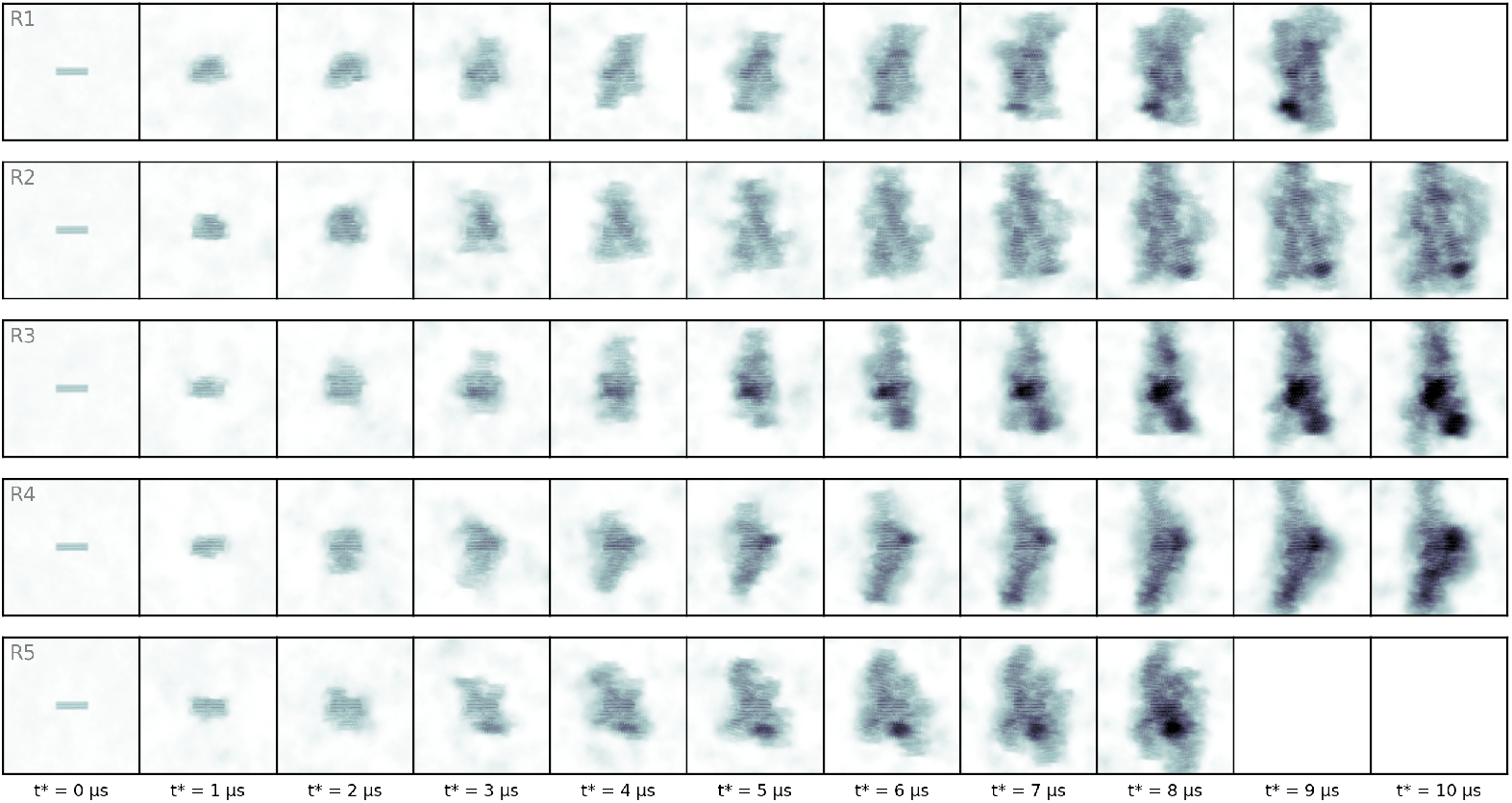
Amyloid growth for five replicas of seeded simulations (polyQ48)

**Suppl. Fig. S13.**
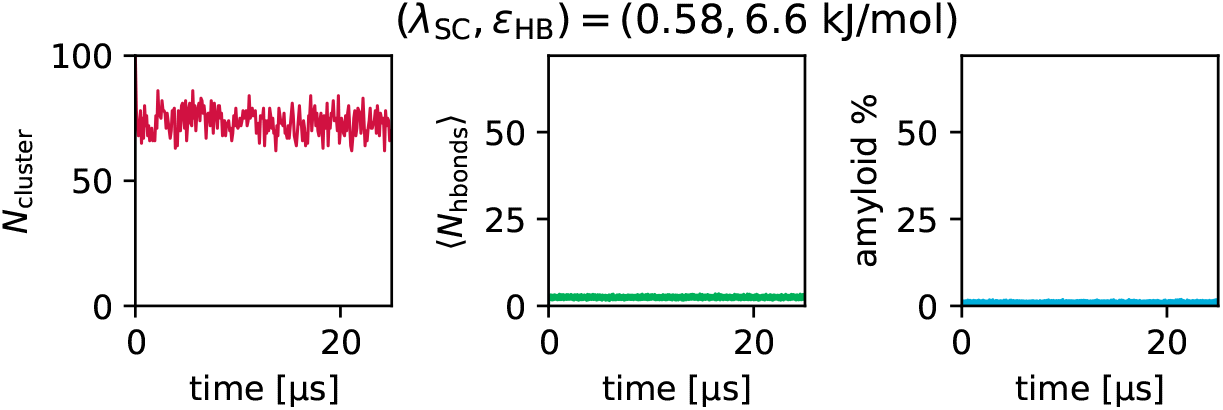
Aggregation simulation analysis for a 25 µs simulation using the default (λ_SC_, *ε*_HB_) values.

**Suppl. Movie S1. Animation of a simulation trajectory depicting amyloid growth through** β**-sheet elongation**. The animation corresponds to the snapshots shown in Fig. 3a–c.

**Suppl. Movie S2. Animation of a simulation trajectory depicting amyloid growth through the zipper mechanism**. The animation corresponds to the snapshots shown in Fig. 3d–f.

**Suppl. Movie S3. Two-dimensional animation of a simulation trajectory from a Q48 seeded simulation**. The animation illustrates the first 5 microseconds of the seeded simulation (replica 3), which is also referenced in Fig. 4e.

